# Small subsets of highly connected taxa predict compositional change in microbial communities

**DOI:** 10.1101/159087

**Authors:** Cristina M. Herren, Katherine D. McMahon

## Abstract

For decades, ecological theory has predicted that the complexity of communities should be related to their stability. However, this prediction has rarely been tested empirically, because of both the difficulty of finding suitable systems where the question is tractable and the trouble of defining “stability” in real systems. Microbial communities provide the opportunity to investigate a related question: how does community connectivity relate to the rate of compositional turnover? We used a newly developed metric called community “cohesion” to test how microbial community connectivity relates to Bray-Curtis dissimilarity through time. In three long-term datasets, we found that stronger connectivity corresponded to lower rates of compositional turnover. Using two case studies of disturbed and reference communities, we found that the predictive power of community connectivity was diminished by external disturbance. Finally, we tested whether the highly connected taxa were disproportionately important in explaining compositional turnover. We found that subsets of highly connected “keystone” taxa, generally comprising 1-5% of community richness, explained community turnover better than using all taxa. Our results suggest that stronger biotic interactions within microbial community dynamics are stabilizing to community composition, and that highly connected taxa are good indicators of pending community shifts.

## Introduction

Theoretical ecologists have studied the relationship between community complexity and stability for decades (MacArthur 1955, May 1972, Pimm 1979, Neutel et al. 2007). Initial results suggesting that complex communities should be unstable (May 1972) prompted a rich literature aimed at understanding how complex communities persist in nature. The primary source of “complexity” considered in these studies is the strength of species interactions (May 2001). These theoretical studies consistently find that connectivity arising from species interactions is a major contributor to community stability (McCann et al. 1998, Ives et al. 2000, Neutel et al. 2002, Wiliams and Martinez 2004). Depending on the configuration and strength of species interactions within a community, greater connectivity can lead to increased or diminished stability (Allesina and Tang 2012).

Despite the substantial theoretical literature on how complexity influences stability, comparatively few studies have investigated this question empirically (but see Kondoh 2008, Neutel and Thorne 2014, Jacquet et al. 2016). This is partly due to the logistical challenges of addressing this question in real systems; such challenges include the difficulty in quantifying species interactions (Laska and Wootton 1998), the need to observe many taxa to satisfy model assumptions (Allesina and Tang 2012), the difficulty of sampling communities completely (Polis 1991), and the need to collect data spanning many generations of the study organisms (Morin and Lawler 1995). Another practical hurdle is defining the terms “complexity” and “stability” for real communities (Connell and Sousa 1983, Neubert and Caswell 1997). Studies that have tested how community complexity relates to community stability have found mixed results. Recently, an analysis of 116 food webs found no consistent pattern between complexity and stability (Jacquet et al. 2016). However, prior studies have found evidence of positive (Polis and Strong 1996, Fagan 1997, Dunne et al. 2002) and negative (Pimm and Lawton 1978, Stouffer and Bascompte 2011) relationships. Thus, relatively little of the ecological theory regarding the complexity-stability debate has been tested empirically, and results of these empirical studies are mixed.

Microbial communities are promising systems for investigating the relationship between community structure and community stability. Several characteristics of microbial communities make it possible to overcome the previously described challenges of testing theoretical hypotheses in empirical systems; microbial communities are sufficiently diverse as to meet the richness assumptions of theoretical models, hundreds of generations can be observed within one dataset, and the resolution of next generation sequencing datasets means that even rare taxa (< 0.01% of communities) are sampled. However, one prominent challenge of testing ecological hypotheses in microbial communities is that interactions between taxa are difficult to observe and therefore must be inferred from observed population dynamics. For this reason, we previously created a robust metric, called “cohesion,” that quantifies the instantaneous connectivity of microbial communities (Herren and McMahon *in press*). Briefly, this method quantifies connectedness values for each taxon in a dataset based on its average correlations with other taxa. Cohesion metrics are calculated from the abundance and connectedness of the taxa present in each community. When many highly connected taxa are present, the cohesion values for a community are larger in magnitude. There are two cohesion values for each sample, corresponding to connectivity arising from positive taxon relationships and connectivity arising from negative taxon relationships.

Recent studies have hypothesized that biotic interactions are important for mediating compositional stability in microbial communities. For example, Zelezniak et al. (2015) found that persistent sub-networks within microbial communities often included a high degree of facilitation among taxa. This result suggested that facilitation reinforces existing community composition, leading to lower rates of compositional change. Furthermore, several microbial studies have found evidence of “keystone taxa,” which are highly interactive and have a disproportionate effect on their communities (Vick-Majors et al. 2014, Agler et al. 2016, Banerjee et al. 2016). Changes in the abundance of keystone taxa lead to shifts in community composition due to cascading effects on other taxa (Mills et al. 1993). Finally, viruses and protists constitute a major source of mortality in marine bacterial communities (Fuhrman and Noble 1995, Suttle 2007), indicating the importance of predation in shaping community composition. Thus, multiple lines of evidence suggest that the strength of biotic interactions within microbial communities should be related to the rate of compositional change.

In this study, we use three long-term microbial datasets (each spanning 10+ years) to test the hypothesis that higher connectivity in microbial communities is related to greater compositional stability through time. As mentioned previously, it is difficult to quantify stability for empirical systems. Instead, we use a related metric as our response variable, which is compositional turnover through time (Bray-Curtis dissimilarity). Modeling Bray-Curtis dissimilarity is a major aim of microbial ecology (Larsen et al. 2012), because the function of microbial communities is expected to change in parallel with changes in community composition (Urich et al. 2008, Sekirov et al. 2010). We also asked whether highly connected “keystone” taxa are disproportionally important for explaining compositional turnover (Power et al. 1996, Jordán et al. 1999). To answer this question, we repeated our analyses of community connectivity versus compositional stability using only the highly connected taxa. Finally, we reasoned that the influence of biotic interactions on compositional change would be diminished when external disturbance to a community was high (Dai et al. 2017). To test this hypothesis, we analyzed two case studies where communities experienced different levels of disturbances. We hypothesized that connectivity would be a better predictor of compositional change when external forcing, and disturbance, was lower. Together, these analyses aimed to identify the conditions under which connectivity is related to compositional turnover and to investigate which taxa are most informative about overall community changes.

## Methods

### Datasets

To test our hypotheses about 1) the relationship between connectivity and compositional turnover and 2) the influence of highly connected taxa, we obtained three long-term, publicly available microbial datasets. These included the San Pedro Ocean Time Series bacterial dataset (SPOT) from the coastal ocean near southern California (described in detail in Cram et al. 2015a), the Lake Mendota (Wisconsin, USA) phytoplankton dataset (ME-phyto, described in detail at https://lter.limnology.wisc.edu), and the Lake Mendota bacterial dataset (ME-bact, described in detail at https://lter.limnology.wisc.edu). Additional information and references for datasets can be found in the Supplementary Online Materials (SOM). We chose these datasets because of their long duration (SPOT: 10 years, ME-phyto: 19 years, ME-bact: 11 years), their large number of samples (SPOT: 274 samples with 437 taxa, ME-phyto: 293 samples with 409 taxa, ME-bact: 91 samples with 7081 taxa), and the variety of technologies used to obtain the datasets (SPOT: automated ribosomal intergenic spacer analysis [ARISA], ME-phyto: cell counts under microscope, ME-bact: 16S rRNA gene amplicon sequencing).

We identified two case studies where comparable microbial communities experienced differing levels of external disturbance. The first case study is the comparison of the phytoplankton communities in Peter Lake and Paul Lake in northern Wisconsin, USA (described in Elser and Carpenter 1988, Cottingham et al. 1998). These lakes were originally one water body, but were artificially divided into two lakes for the purpose of conducting ecological disturbance experiments. Paul Lake served as the undisturbed reference system, while Peter Lake was experimentally disturbed using nutrient supplementation and fish additions over the course of the time series (see SOM). Each lake was sampled 197 times over 12 years. Phytoplankton taxa were enumerated using direct cell counts under a microscope.

The second disturbance case study is a comparison between two types of plaque communities sampled as a part of the Human Microbiome Project (HMP). Briefly, samples were collected from 242 human volunteers at up to 18 body sites at two sample collection dates with a maximum interval of 14 days. We compared the bacterial communities from the highly disturbed, exposed plaque site (supragingival plaque) to the protected plaque site beneath the gums (subgingival plaque). For both sites, we evaluated the relationship between community cohesion and compositional turnover (Bray-Curtis dissimilarity) in an individual’s microbiome between the two sampling times.

### Hypotheses: Long-Term Datasets

Following the result that persistent microbial sub-networks are enriched in taxon interactions (Zelezniak et al. 2015), we expected that greater connectivity would be related to lower compositional change. Additionally, we hypothesized that the highly connected taxa would have a disproportionate influence on community dynamics. Thus, we expected that subsets of highly connected taxa would be better predictors of community turnover (Bray-Curtis dissimilarity) than randomly chosen subsets of taxa.

### Hypotheses: Case Study Comparisons

For Peter Lake and Paul Lake, we reasoned that the experimental perturbations would be a cause of community composition change in Peter Lake, but not in the undisturbed Paul Lake. Therefore, we expected that biotic interactions would contribute less to compositional turnover in the disturbed lake, Peter Lake. Thus, we hypothesized that community cohesion would be a better predictor of Bray-Curtis dissimilarity in the undisturbed Paul Lake than in Peter Lake.

For the two plaque bacterial communities, we reasoned that compositional change at the exposed site (supragingival plaque) would be influenced more strongly by immigration and dispersal than by biotic interactions. Conversely, we expected that the protected plaque communities (subgingval plaque) would be influenced by biotic interactions, because taxa are contained in close proximity for long periods of time. Thus, we hypothesized that cohesion would be a significant predictor of Bray-Curtis dissimilarity for the protected plaque site (subgingival plaque), but not at the exposed site (supragingival plaque).

### Statistical Methods

We used cohesion metrics (Herren and McMahon *in press*) as a measure of the connectivity of the microbial communities (see SOM). We calculated cohesion metrics for the five datasets (three long-term time series and two case studies). Briefly, this workflow calculates two metrics for each sample quantifying the connectivity due to positive correlations between taxa and connectivity due to negative correlations between taxa. Cohesion metrics are calculated for each sample by taking the sum of every taxon’s connectedness score (also calculated within the cohesion workflow) multiplied by its abundance in the sample.

For each dataset, we conducted linear regressions modeling the compositional turnover (Bray-Curtis dissimilarity) between time points as a function of the cohesion metrics. Stated another way, we asked whether cohesion metrics predict Bray-Curtis dissimilarity. For the SPOT dataset, we analyzed the bacterial communities from the chlorophyll maximum site, reasoning that the cholorophyll maximum site represented a discrete ecological community. For taxa in the HMP plaque datasets, we calculated taxon connectedness values using correlations between taxa among individuals at the first sampling timepoint. Additional methods and the parameter values used in the workflow for each dataset can be found in Supplementary Table 1.

To test the hypothesis that highly connected taxa are disproportionately influential in determining community dynamics, we iteratively repeated the regression analysis (modeling Bray-Curtis dissimilarity as a function of community cohesion), each time calculating cohesion from different subsets of taxa. We excluded taxa based on their connectedness values, where we removed the least connected taxa first. For example, when 40 taxa were included in the analysis, the negative cohesion metric was calculated from the 40 taxa with the strongest negative connectedness, and the positive cohesion metric was calculated from the 40 taxa with the strongest positive connectedness. We recorded the R^2^ value from the linear model (Bray-Curtis dissimilarity vs. cohesion) for each subset of taxa.

We then repeated the workflow described above (removing taxa and running the linear regression) using random subsets of taxa, rather than using the most highly connected taxa. Thus, when 40 taxa were included, we randomly selected 40 taxa from which to calculate the positive and negative cohesion values. We recorded the model R^2^ value of the linear regression when taxa were randomly included in the workflow. Then, we repeated this process 500 times, as to generate a distribution of model R^2^ values when 40 random taxa were selected. We ran 500 models for each possible number of taxa included in the workflow. We had hypothesized that the highly connected taxa would be more informative about overall community changes than randomly chosen taxa; thus, we expected that the model using the highly connected taxa would have a larger R^2^ value.

## Results

### Long-Term Datasets

For each of the three long-term datasets (SPOT, ME-phyto, and ME-bact), we used linear regression to analyze the amount of variability in community composition turnover (Bray-Curtis Dissimilarity) that could be explained by community connectivity (cohesion metrics). Representative results from all datasets analyzed are presented in Table 1.

**Table 1:** Representative Results of Cohesion as a Predictor of Bray-Curtis Dissimilarity

| *Dataset* | *Total # of Taxa* | *Optimal # of Taxa\** | *Maximum Adjusted $R^2$* | *Positive Cohesion P value* | *Negative Cohesion P value* | *Positive Cohesion Direction <sup>+</sup>* | *Negative Cohesion Direction <sup>+</sup>* | *Data Points in Analysis <sup>#</sup>* |
| --- | --- | --- | --- | --- | --- | --- | --- | --- |
| ME - phyto | 409 | 15 | 0.485 | n.s. | $< 1*10^{-27}$ | NA | Stronger is stabilizing | 186 |
| ME - bact | 7081 | 33 | 0.428 | $< 1*10^{-3}$ | $< 1*10^{-7}$ | Stronger is stabilizing | Stronger is stabilizing | 54 |
| SPOT – Chl. Max. | 392 | 15 | 0.478 | n.s. | $< 1*10^{-4}$ | NA | Stronger is stabilizing | 36 |
| Protected Plaque | 2190 | 13 | 0.207 | $< 1*10^{-4}$ | 0.014 | Stronger is stabilizing | Weaker is stabilizing | 93 |
| Exposed Plaque | 2124 | 79 | 0.018 | n.s. | n.s. | NA | NA | 95 |
| Paul - Reference | 209 | 13 | 0.487 | $< 1*10^{-5}$ | $< 1*10^{-18}$ | Weaker is stabilizing | Stronger is stabilizing | 123 |
| Peter - Disturbed | 237 | 57 | 0.374 | 0.009 | $< 1*10^{-6}$ | Stronger is stabilizing | Stronger is stabilizing | 121 |

\* Indicates the number of taxa where the maximum adjusted $R^2$ value occurred
+ These columns indicate the direction of a significant relationship between cohesion and Bray-Curtis dissimilarity. For example, “stronger is stabilizing” means that greater cohesion is related to lower Bray-Curtis dissimilarity. Non-significant relationships are denoted “n.s.”.

Stronger cohesion, whether positive or negative, was consistently and significantly related to lower rates of compositional change (Table 1). Stronger negative cohesion was significantly related to lower Bray-Curtis dissimilarity in all three datasets (Fig. 1B, D, F). In the ME-bact dataset, stronger positive cohesion was also significantly related to lower compositional turnover (Table 1). Maximum adjusted model R^2^ values were 0.485 for ME-phyto, 0.428 for ME-bact, and 0.478 for SPOT chlorophyll maximum.

**Figure 1:**
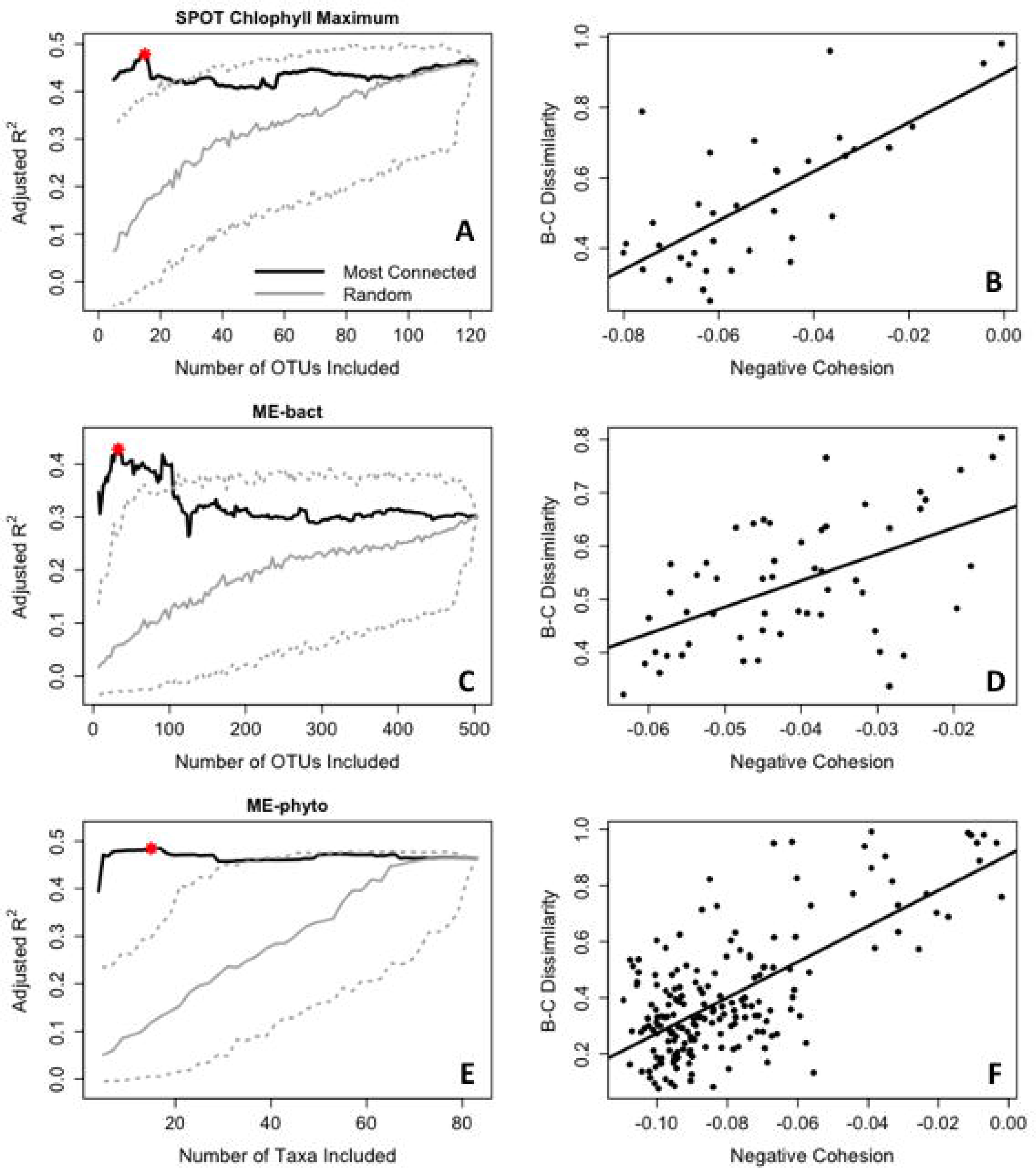
Analyses of the three long-term microbial datasets show that stronger cohesion is related to lower compositional turnover in all three long-term datasets. Left-hand panels (A, C, E) show the how the adjusted model R^2^ values of the regression analysis changed as taxa were excluded from cohesion calculations. For each number of taxa on the x-axis, cohesion values were calculated from the most highly connected taxa (black line) and from a random subset of taxa (grey lines). The solid grey line shows the median adjusted model R^2^ for randomly selected subsets, while the dashed grey lines give the 5% and 95% intervals. Median R^2^ values from models using random subsets of taxa declined as fewer taxa were included in the cohesion metrics. When 1-5% of taxa within a community were used to calculate cohesion, models using highly connected taxa generally had higher model R^2^ values than models using random taxa. The red stars in left-hand panels identify the regression model with the highest adjusted R^2^, which is displayed in the paired right-hand panel. Right-hand panels (A, C, E) show the best-fitting linear regressions modeling compositional turnover (Bray-Curtis dissimilarity) as a function of cohesion from negative connections between taxa. Points indicate Bray-Curtis dissimilarity between sequential samples. Solid lines show the fit of linear models. All three datasets showed that cohesion arising from negative correlations between taxa was a strong predictor of Bray-Curtis dissimilarity (Table 1).

We re-calculated cohesion metrics from subsets of highly connected taxa in order to evaluate whether highly connected taxa were disproportionately informative about compositional turnover (black line, Fig. 1A, C, E). We also calculated cohesion metrics using random subsets of taxa to evaluate whether highly connected taxa modeled Bray-Curtis dissimilarity better than randomly chosen taxa (grey lines, Fig. 1A, C, E). In the models containing random subsets of taxa, model R^2^ values declined as fewer taxa were included in cohesion calculations (solid grey line indicates the median). Conversely, in models using the most highly connected taxa, the adjusted R^2^ values remained stable as the least-connected taxa were removed (black line). In all three long-term datasets, adjusted R^2^ values increased when small subsets (< 5% total richness) of highly connected taxa were included (Table 1). Maximum R^2^ values occurred when using 15 taxa in ME-phyto, 33 taxa in ME-bact, and 15 taxa in SPOT (Fig. 1A, C, E).

In all three datasets, models based on the most highly connected taxa to calculate cohesion significantly outperformed the models using random subsets of taxa when small proportions of taxa were included. Significance was determined as instances when the model R^2^ value using highly connected taxa was above the 95^th^ percentile of R^2^ values from models using random taxa. For the SPOT dataset, the model using highly connected taxa performed significantly better than the model using randomly selected taxa when fewer than 25 taxa were included. For the ME-phyto dataset, it was when fewer than 35 taxa were included. For the ME-bact dataset, it was fewer than 105 taxa.

### Identities of Highly Connected Taxa

We were curious about the identities of the most highly connected taxa in the three long-term datasets. We focused on taxa that had the strongest negative associations with other taxa, because negative cohesion was highly significant in all long-term datasets (Fig. 1, Table 1). In the ME-phyto dataset, eight of the ten taxa with the largest negative connectedness values were cyanobacteria (see SOM for list). For the ME-bact dataset, we compared the lists of the fifty most abundant taxa and the fifty taxa with largest negative connectedness values (see SOM). Twenty-two taxa were on both these lists. Twenty-eight taxa were among the fifty most connected but not the fifty most abundant. These included three of the four recognized clades in the acIV Actinobacteria lineage, a member of the Chloroflexi phylum, and two members of the Planctomycetes phylum, all of which are relatively understudied by freshwater microbial ecologists. Among the Proteobacteria in this list were PnecD, a relatively rare member of the genus *Polynucleobacter*, and several members of the order Rhizobiales. Although these organisms are not among the most ubiquitous or abundant taxa found in freshwater lakes, the results obtained here motivate us to study their ecology more intently, particularly with genome-based methods.

### Case Study: Peter Lake and Paul Lake

As with the long-term datasets, we used cohesion metrics as predictors of Bray-Curtis dissimilarity for phytoplankton communities in Peter and Paul Lakes. We had hypothesized that cohesion metrics would be better predictors of compositional change in the reference system, Paul Lake. We conducted separate analyses for the two lakes.

As expected, cohesion metrics were better predictors for Paul Lake than for the disturbed system, Peter Lake. We evaluated this prediction by comparing model R^2^ values for the two lakes (Fig. 2). Across nearly the entire range of taxa included, models analyzing the Bray-Curtis dissimilarity of phytoplankton communities in Paul Lake had a higher R^2^ value than similar models for Peter Lake. The exception was when very few (< 10) taxa were included in the cohesion calculations. In both lakes, model R^2^ values dropped significantly when fewer than 10 taxa were used to calculate cohesion. The best model fit in Paul Lake occurred when 13 taxa were included (adjusted R^2^ = 0.487), whereas for Peter Lake it was 57 taxa (adjusted R^2^ = 0.374). In Paul Lake, stronger negative cohesion and weaker positive cohesion was both significantly related to lower Bray-Curtis dissimilarity (Table 1). In Peter Lake, stronger negative cohesion and stronger positive cohesion were both significantly related to lower Bray-Curtis dissimilarity (Table 1).

**Figure 2:**
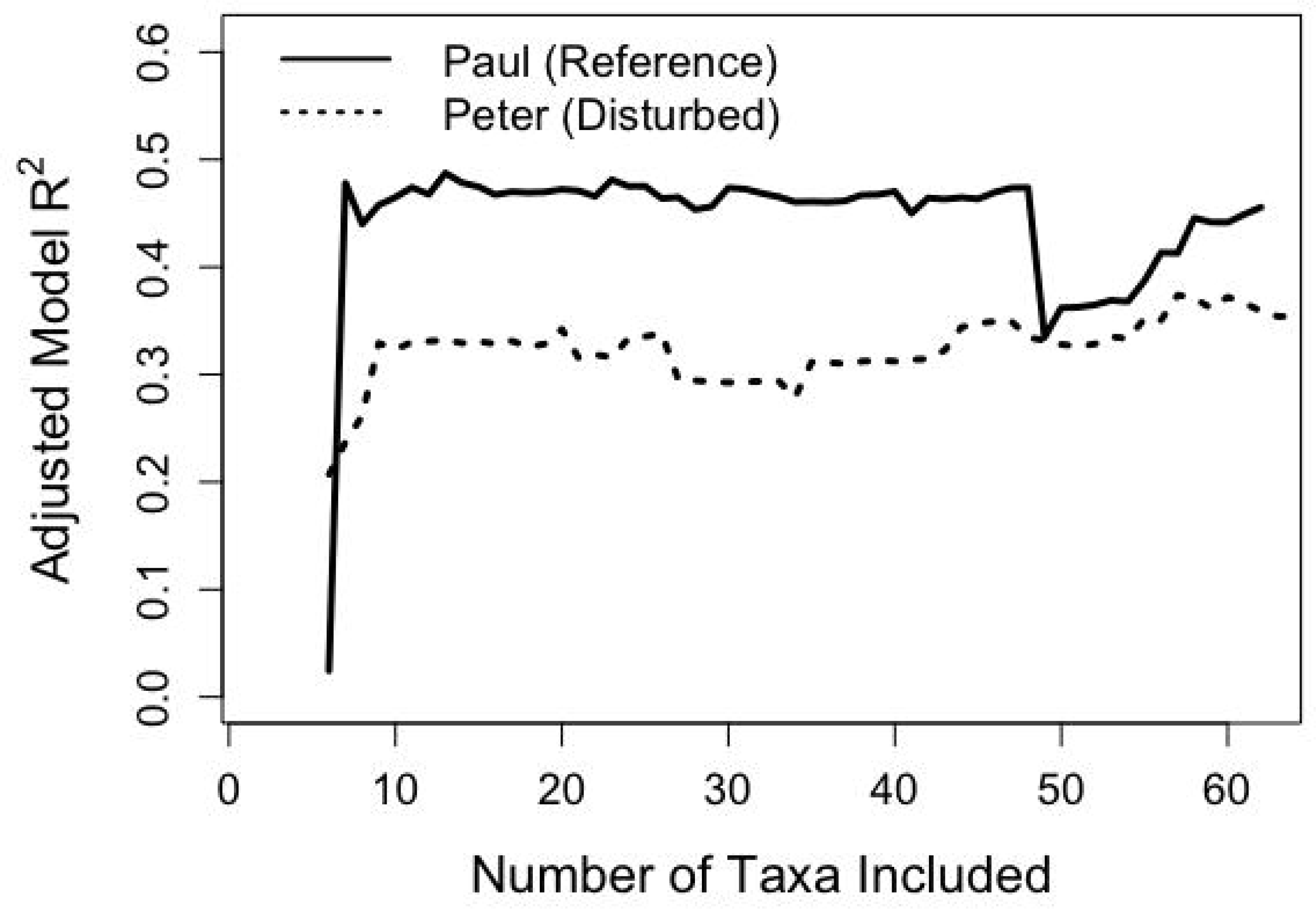
Cohesion explained a greater amount of variability in phytoplankton community turnover in the undisturbed Paul Lake, as compared to an experimentally disturbed system, Peter Lake. The model R^2^ values predicting Bray-Curtis dissimilarity in Paul Lake were generally higher than for models predicting Bray-Curtis dissimilarity in Peter Lake. The exception was when models used very few (< 10) taxa to calculate cohesion metrics. As in Figure 1, taxa were sequentially removed from the analysis in reverse order of their connectedness (i.e. least connected taxon removed first).

### Case Study: Exposed and Protected Plaque Communities

We tested whether cohesion could explain community composition turnover in plaque communities in the human-associated microbiome. We expected that cohesion would be a significant predictor of compositional turnover at the protected plaque site (subgingival plaque), but not at the exposed plaque site (supragingival plaque). In this analysis, we calculated Bray-Curtis dissimilarities from two communities sampled from the same individual host, collected at two different time points.

In the exposed plaque communities (supragingival plaque), we found that there was no significant relationship between either cohesion metric and Bray-Curtis dissimilarity (Fig. 3). However, in the protected plaque communities (subgingival plaque), cohesion was significantly related to Bray-Curtis dissimilarity. The model fit was best (adjusted R^2^ = 0.207) when 13 OTUs were included (Fig. 3). Stronger positive cohesion and weaker negative cohesion were both significantly related to lower Bray-Curtis dissimilarity (Table 1).

**Figure 3:**
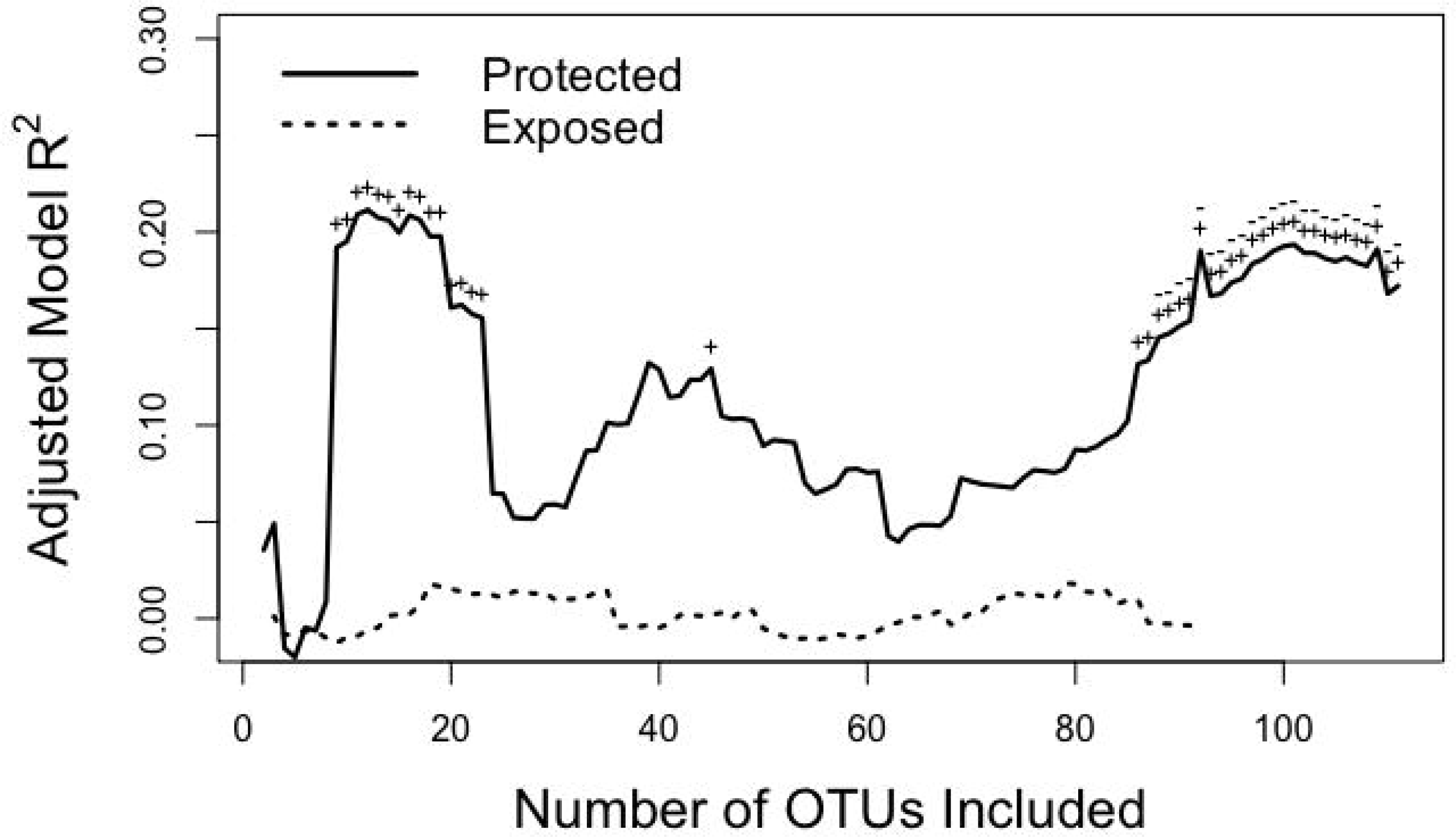
The adjusted model R^2^ values for plaque communities sampled as part of the Human Microbiome Project show that cohesion was a significant predictor of Bray-Curtis dissimilarity in the protected plaque site (subgingival plaque, solid line), but not at the exposed plaque site (supragingival plaque, dashed line). Icons above the solid line indicate when positive cohesion was significant at p < 0.001 (+) and when negative cohesion was significant at p < 0.001 (-). At the exposed plaque site, cohesion was never a significant (p < 0.05) predictor of Bray-Curtis dissimilarity.

We conducted this same analysis using the other 16 body sites sampled as a part of the Human Microbiome Project (see SOM for results). Most sites (11 of 16) showed highly significant relationships (p < 0.001) between cohesion and the rate of compositional turnover (Bray-Curtis dissimilarity). At all 11 sites, stronger negative cohesion was related to lower Bray-Curtis dissimilarity. Positive cohesion was highly significant at 6 of the 11 sites, but showed mixed relationships with Bray-Curtis dissimilarity.

## Discussion

The consistent results from the three long-term (10+ year) microbial time series showed that stronger connectivity within aquatic microbial communities was related to greater compositional stability. In all three cases, stronger cohesion values were significantly related to lower Bray-Curtis dissimilarity over time (Fig. 1B, D, F). Moreover, models using information from small subsets of highly connected taxa predicted compositional turnover performed better than models using all taxa (Fig. 1A, C, E). Therefore, the most highly connected taxa had the strongest relationship with compositional change, and their presence corresponded to increased compositional stability. In all three long-term datasets, highly connected taxa performed significantly better than models built using random assemblages of taxa. Only a small fraction of taxa, generally comprising 1-5% of total richness, were necessary to model compositional turnover. These qualitatively consistent results show support for the hypotheses that 1) community connectivity is a strong mediator of compositional stability and 2) highly connected taxa have disproportionate influence on observed community dynamics.

The predictive power of our models in the long-term datasets was striking, given that no environmental factors were included in these analyses. For the three long-term datasets, the model R^2^ values ranged between 0.4 and 0.5. For comparison, previous analyses modeling the community similarity between time points in the SPOT dataset obtained maximum R^2^ values of approximately 0.2, even when using over 30 environmental parameters (Cram et al. 2015a). Similarly, a model explaining compositional turnover in the ME-phyto dataset using environmental variables had an adjusted R^2^ value of 0.23 (Herren and McMahon *in press*).

Our result that stronger negative cohesion was related to lower compositional turnover in the long-term time series was consistent across a variety of ecosystems, sampling methods, and sample dates. The three datasets were obtained using different techniques for determining abundance, including direct cell counts (ME-phyto), 16S rRNA gene tag sequencing (ME-bact), and ARISA (SPOT). These methods all differ in their sensitivity and bias. Thus, the consistency of our results suggest that including cohesion as a predictor variable might improve models of compositional turnover in many microbial systems.

### Disturbance Decreases the Importance of Connectivity

The case studies of disturbed systems showed that community cohesion had less explanatory power when communities experienced external disturbance. The Peter Lake vs. Paul Lake comparison showed that cohesion metrics were better predictors of Bray-Curtis dissimilarity in the undisturbed system, Paul Lake (Fig. 2). In Peter Lake, experimental perturbations caused shifts in the phytoplankton community (Carpenter et al. 1987, Carpenter et al. 1996, Cottingham and Carpenter 1998). Thus, some of the compositional change in Peter Lake was due to experimental disturbances. Our results agree with these previous conclusions, suggesting that connectivity had decreased influence on compositional change in the perturbed lake, Peter Lake.

Analyses of the protected and exposed plaque sites showed that community cohesion was only an important explanatory factor in compositional turnover at the protected plaque site (Fig. 3). Many of the same OTUs were present in the protected and exposed plaque communities, but their connectedness and power to predict compositional change were different at the two sites. These results suggest that high levels of disturbance and dispersal can disrupt the relationship between biotic interactions and community stability.

There are two main ways in which disturbance can alter the relationship between biotic interactions and compositional change. First, disturbances causing high immigration or emigration of taxa disrupt established species interactions. Biotic interactions drive population dynamics by influencing taxon growth and death rates (Gotelli 2001); thus, the effects of biotic interactions will be most apparent when taxa interact consistently over many generations. Second, disturbances cause compositional change that is not linked to biotic interactions. For example, compositional change at the exposed plaque site may have resulted from tooth brushing or from consuming food. Thus, the proportion of total compositional change due to biotic interactions would be diminished in this case. The lower predictive power of cohesion when applied to highly disturbed communities suggests that the importance of biotic interactions in community assembly and turnover is context dependent.

### Highly Connected Taxa as Keystone Taxa

Focusing on highly connected taxa may be a useful strategy for researchers seeking to understand microbial community assembly and compositional change. In all three long-term datasets, the ability to explain compositional turnover was highest when a small number (15-33) of highly connected taxa were included. Similarly, in the two reference systems in the case study analyses, the optimal number of taxa to include was 13 for both datasets (Table 1). Including taxa with weaker connectedness values in our models often obscured the signal of connectivity captured by the cohesion metrics. These results support the hypothesis that the highly connected taxa may function as “keystone taxa” within microbial communities; the relatively small subsets of highly connected taxa had outsized explanatory power of overall community dynamics. Additionally, although some of the datasets contained the same phytoplankton taxa (ME-phyto, Peter Lake, Paul Lake), the same taxon received different scores of connectedness in the various datasets. This result suggests that the ecological context of the microbial communities is important for determining which taxa will act as keystone taxa in various environments.

We propose that the approach of evaluating model fit using different subsets of taxa could be generalized to other analyses with different response variables. Model fit should be best when the most informative taxa are included in the analysis. One strategy for identifying taxa with disproportionate influence would be to include the taxa where the model R^2^ values spike in Fig. 1. Although model R^2^ values remained high when small numbers of taxa were included, we would caution against building predictive models with fewer than 5-10 taxa. In this case, cohesion values obtained from a training set of communities may be prone to high variability when applied to new communities, especially if there are directional trends in taxon abundances over time.

### Ecological Interpretation of Connectivity and Compositional Turnover

In the majority of instances where cohesion metrics were significant predictors of Bray-Curtis dissimilarity, stronger connectivity was related to greater compositional stability. However, there were cases that deviated from this norm, where stronger connectivity was related to more rapid change. We hypothesize that these anomalies are mediated by the ecology of the different study sites. For example, the result from Paul Lake that stronger positive cohesion was destabilizing was driven by samples from the summer of 1993, when a large and persistent cyanobacterial bloom disrupted normal seasonal dynamics. Similarly, following the result that cohesion had lower explanatory power in disturbed systems, it would be interesting to investigate how the strength of deterministic versus stochastic forces alters the relationship between community connectivity and community stability. This might be done with the Human Microbiome Project dataset, as immigration and selective pressure likely differ between body sites (Li and Ma 2016). Our preliminary analysis of this dataset showed that 12 of the 18 sites had a strong relationship between cohesion values and compositional turnover rate, but the explanatory power of the models varied. Quantifying dispersal and selection rates at different sites may shed light on the variability of the observed relationships and the degree to which community connectivity can explain compositional change.

Under the assumption that cohesion measures biotic interactions (Herren and McMahon *in press*), our results support the hypothesis that biotic interactions are stabilizing to microbial community composition. Several recent studies have concluded that biotic interactions can be strong drivers of microbial population dynamics, on par with or exceeding the influence of environmental factors (Cram et al. 2015b, Lima-Mendez et al. 2015, Weitz et al. 2016, Cabello et al. 2016, Trivedi et al. 2017). For example, many OTUs are more strongly related to other OTUs than to habitat variables (Cram et al. 2015b). However, few studies have tested the relationship between connectivity and compositional change, primarily because the methods to quantify connectivity have only been recently developed. Initial theoretical studies indicated that stronger biotic linkages would be destabilizing to ecological communities (May 1972, Pimm 1979). However, these initial studies also made several simplifying assumptions about the organization of ecological food webs. The ensuing literature has discovered several possible mechanisms that allow diverse and complex communities to persist in nature (e.g. McCann et al. 1998, Brose et al. 2006, Kondoh 2006). Future work might consider how the attributes of microbial communities, including spatial structuring (Long and Azam 2001), dispersal rates (Finlay 2002), and the possibility of dormancy (Lennon and Jones 2011) influence the relationship between connectivity and compositional stability.

Biotic interactions create feedback loops within ecological communities that can amplify or dampen the effects of external perturbations (Berryman and Millstein 1989). One mechanistic hypothesis for the result that stronger connectivity is related to lower compositional change is that the taxon interactions in microbial communities are arranged to form negative feedback loops, thereby mitigating the effects of disturbance (Konopka et al. 2015, Coyte et al. 2015). Thus, stronger interactions would lead to stronger negative feedback loops that buffer communities from compositional change. Our findings also agree with recent work showing that persistent modules of taxa are enriched in taxon interactions (Zelezniak et al. 2015). Thus, another interpretation is that biotic interactions create self-reinforcing modules within bacterial communities, which leads to lower turnover. One possible mechanism generating these self-reinforcing subunits is metabolite exchange between taxa (Morris et al. 2013, Levy and Borenstein 2013). Finally, our work agrees with recent studies that hypothesize that microbial communities contain keystone taxa, which shape community assembly due to their strong interactions with other taxa (Vick-Majors et al. 2014, Agler et al. 2016, Banerjee et al. 2016). We propose that studying these keystone taxa might allow researchers to prioritize organism-centric studies to learn why and how specific taxa have such a strong influence on communities.

Ecological theory offers some insight into why negative cohesion was often more strongly related to compositional stability than positive cohesion. Under some circumstances, pairwise correlations may be indicative of pairwise taxon interactions; we make this simplifying assumption to investigate our results in the context of classical ecological theory. Mathematical models using local stability analysis with simple communities have indicated that stable equilibria are common when negative interactions (e.g. competition, predation) are present. For instance, scenarios with stable eqiulibria include: two or more competitors (May and Leonard 1975), one predator and one prey (Rosenzweig and MacArthur 1963), one predator with multiple prey (Holt 1977), and multiple predators with one or more prey (McPeek 2012). Conversely, stability is rare in food webs with exclusively positive pairwise interactions (May 1981). However, recent theoretical literature has indicated that mutualism within the context of other, negative interactions can be stabilizing (Mougi and Kondoh 2012). Thus, ecological theory indicates that the placement and strength of negative interactions within communities is critical to maintaining stable composition. The traits of the most highly connected Mendota phytoplankton taxa further support this line of reasoning. Most of the taxa associated with low compositional turnover were cyanobacteria, which often have a competitive relationship with other phytoplankton (Fong et al. 1993). Several studies have documented the self-reinforcing effect of competition for light in aquatic environments, showing that high cyanobacterial abundance can be a stable state in eutrophic lakes (Scheffer et al. 1997, Schröder et al. 2005). Thus, our results align with existing theoretical explanations of phytoplankton community transitions, and suggest that similar dynamics may be present in other systems. Although the knowledge of traits and interactions is scarce for taxa in the other two long-term datasets (SPOT and Mendota-bact), the lists of highly connected taxa provided here (available in SOM) may be useful starting places for trait-based studies.

We encourage future studies to examine traits of highly connected taxa using modeling or experimental approaches. Although the assumption that taxon correlations are indicative of taxon interactions is useful for invoking ecological theory, there are several conditions where this assumption would be false. For example, two competing taxa might show a negative correlation in their abundances through time due to competitive exclusion; conversely, two competing taxa may have similar niches, and therefore might show a positive correlation due to simultaneous responses to environmental drivers. Finally, other trophic levels likely influence the correlations and connectedness metrics observed in these microbial communities, although these factors are not explicitly included in these analyses. Thus, mechanistic models would greatly benefit further studies of the role of highly connected taxa in community dynamics.

Our results show several empirical instances where stronger connectivity is related to greater compositional stability, contrary to the initial theoretical finding that highly connected communities should be unstable. Empirical food webs have many non-random attributes, which may explain why our results differ from theoretical expectations and analyses of simulated datasets (Pimm 1980, Polis 1991, Neutel et al. 2007). Our consistent finding that greater community connectivity (especially from negative connections between taxa) results in lower compositional turnover suggests that either evolutionary or community assembly processes arrange biotic interactions to form stabilizing feedback loops.

## Acknowledgements

We thank the owners of the long-term time series for making their datasets publicly available. Data owners and curators include: the Fuhrman lab and the Wrigley Institute at the University of Southern California for the SPOT dataset, the North Temperate Lakes Long Term Ecological Research program for the Lake Mendota phytoplankton and Lake Mendota bacteria datasets, past and present members of the McMahon lab (but especially Ryan Newton, Georgia Wolfe, Emily Read, and Robin Rohwer) and the Earth Microbiome Project for the Lake Mendota bacteria dataset, and the Cascade research group for the Peter Lake and Paul Lake datasets. We thank the Human Microbiome Project for access to their data. Mark McPeek and members of the McMahon lab contributed helpful comments on this work. Finally, we personally thank the individual program directors and leadership at the National Science Foundation for their commitment to continued support of long-term ecological research. This work was funded by a United States National Science Foundation (NSF) GRFP award to CMH (DGE-1256259). KDM acknowledges funding from the NSF Long Term Ecological Research program (NTL-LTER DEB-1440297) and an INSPIRE award (DEB-1344254). This material is also based upon work that supported by the National Institute of Food and Agriculture, U.S. Department of Agriculture (Hatch Project 1002996).

Supplementary material is available at ISME Journal’s website.

